# Global remapping in granule cells and mossy cells of the mouse dentate gyrus

**DOI:** 10.1101/2022.09.06.506812

**Authors:** Sang Hoon Kim, Douglas GoodSmith, Stephanie J. Temme, Fumika Moriya, Guo-li Ming, Kimberly M. Christian, Hongjun Song, James J. Knierim

## Abstract

Hippocampal place cells exhibit spatially modulated firing, or place fields, which can remap to encode changes in the environment or other variables. Unique among hippocampal subregions, the dentate gyrus (DG) has two excitatory populations of place cells, granule cells and mossy cells, which are among the least and most active spatially modulated cells in the hippocampus, respectively. Previous studies of remapping in the DG have drawn different conclusions about whether granule cells exhibit global remapping and contribute to the encoding of context specificity. By recording granule cells and mossy cells as mice foraged in different environments, we found that by most measures, both granule cells and mossy cells remapped robustly but through different mechanisms that are consistent with firing properties of each cell type. Our results resolve the ambiguity surrounding remapping in the DG and suggest that most spatially modulated granule cells contribute to orthogonal representations of distinct spatial contexts.

## INTRODUCTION

Place cells in the hippocampus fire at specific locations when an animal is exploring an environment, resulting in spatially modulated firing known as place fields (O’Keefe, 1976). Place cell activity is thought to provide a foundation for an internal cognitive map to support episodic memory and spatial navigation (O’Keefe and Dostrovsky, 1971; O’Keefe and Nadel, 1978).

These place field maps can reorganize in different environments and can be modulated by spatial and nonspatial changes within a single environment (Bostock et al., 1991; Cressant et al., 2002; Leutgeb et al., 2005; Muller and Kubie, 1987). This remapping process is thought to contribute to episodic memory by supporting context specificity and preventing interference between related memories (Colgin et al., 2008; Knierim, 2003; Nadel and Willner, 1980). Global remapping reflects a statistically independent spatial map of each environment, such that place cells can have a place field in one environment but not another, or place cells that have place fields in both environments are active at independent spatial locations. Some degree of remapping by place cells has been observed in all hippocampal subregions (Lee et al., 2004; Leutgeb et al., 2005; Leutgeb et al., 2004; Lu et al., 2015; Vazdarjanova and Guzowski, 2004; Wills et al., 2005; Wilson and McNaughton, 1993). Remapping in the dentate gyrus (DG), however, is currently a subject of debate as recent studies have made conflicting claims about the extent of remapping by different principal cell types following environmental manipulations (Allegra et al., 2020; GoodSmith et al., 2017; Hainmueller and Bartos, 2018; Senzai and Buzsaki, 2017). Granule cells, in particular, have been variously described as being highly stable with minimal remapping (Hainmueller and Bartos, 2018); showing only “modest” remapping compared to robust remapping of mossy cells (Senzai and Buzsaki, 2017); or exhibiting robust remapping in distinct (GoodSmith et al., 2017) and similar (Allegra et al., 2020) environments.

Aside from differences in experimental design, each study used different measures and definitions of remapping and different inclusion/exclusion criteria for analysis of individual cells. However, the data presented in these papers appear qualitatively similar, raising the question of how such disparate conclusions were reached.

Unlike other regions of the hippocampus where place cells are represented by the sole excitatory population of pyramidal cells, the DG contains two excitatory cell types with place cell properties: granule cells (GCs) and mossy cells (MCs). Relative to pyramidal cells in CA1 and CA3, it has been difficult to probe the electrophysiological signatures of granule cells and mossy cells in freely moving animals due to limited methods for cell-type identification from extracellular DG recordings. Recently, however, a number of in vivo electrophysiological and imaging investigations of the DG have revealed differences in the firing properties of these two populations (Allegra et al., 2020; Danielson et al., 2016; Danielson et al., 2017; Diamantaki et al., 2016; GoodSmith et al., 2017; GoodSmith et al., 2019; Hainmueller and Bartos, 2018; Jung et al., 2019; Jung and McNaughton, 1993; Leutgeb et al., 2007; Senzai and Buzsaki, 2017).

Mossy cells are among the most active excitatory cell type in the hippocampus and frequently exhibit multiple place fields within a single environment (Danielson et al., 2017; GoodSmith et al., 2017; Neunuebel and Knierim, 2012; Senzai and Buzsaki, 2017). Granule cells, on the other hand, rarely fire during exploratory behavior in an open arena and, if so, they typically display a single place field (Diamantaki et al., 2016; GoodSmith et al., 2017; Neunuebel and Knierim, 2012; Senzai and Buzsaki, 2017). This sparse firing within the dense granule cell population is thought to contribute to the segregation of similar inputs through distributed patterns of activation in largely nonoverlapping populations (Diamantaki et al., 2016; Jung and McNaughton, 1993; McNaughton and Morris, 1987; McNaughton and Nadel, 1990). However, recent reports suggest that granule cells exhibit a lower degree of remapping than mossy cells or CA pyramidal cells, arguing against the hypothesized role of granule cells in generating highly orthogonalized representations of similar inputs (Hainmueller and Bartos, 2018; Senzai and Buzsaki, 2017). Resolving the ambiguity surrounding cell type-specific remapping properties is critical to understand how the DG contributes to spatial encoding and discrimination.

To determine the potential source(s) of the conflicting interpretations in previous studies, we performed tetrode recording in the DG from 35 adult mice as they foraged for food in four distinct environments, two of which had the same geometrical shape. We also investigated the reproducibility of granule cell and mossy cell spatial firing patterns by re-exposing mice to one of the four environments at the end of each recording day. By most measures, granule cells and mossy cells exhibited a similar, robust degree of global remapping across environments and maintained a similar degree of correlation between spatial representations during repeated exposures to the same environment. Although granule cells and mossy cells remapped to a similar degree overall, the appearance of global remapping was distinct for each cell type: sparsely firing granule cells remapped through largely nonoverlapping populations of active cells, whereas the highly active mossy cells exhibited a combination of population remapping and spatial decorrelation of place fields across environments.

## RESULTS

### Classification of granule cells and mossy cells

We recorded 420 well-isolated, putative excitatory cells from the DG granule cell layer and hilus of 35 male and female mice as they foraged for food in four distinct environments (Figure S1A). The environments consisted of two squares, one circle and one octagon. Each environment had a unique color and texture on the floors and a unique color and cues on the walls (Figure 1A and S1B). All environments were placed at the same fixed location in a single room. To evaluate the reproducibility of neural activity patterns, the final session of the day was a repeated exposure to one of the four environments. We also recorded neural activity during baseline sleep/rest sessions that preceded and followed the behavior sessions to confirm the stability of recordings. No major, qualitative differences were detected between males and females (Table S1), so the data were pooled together for analysis.

**Figure 1.**
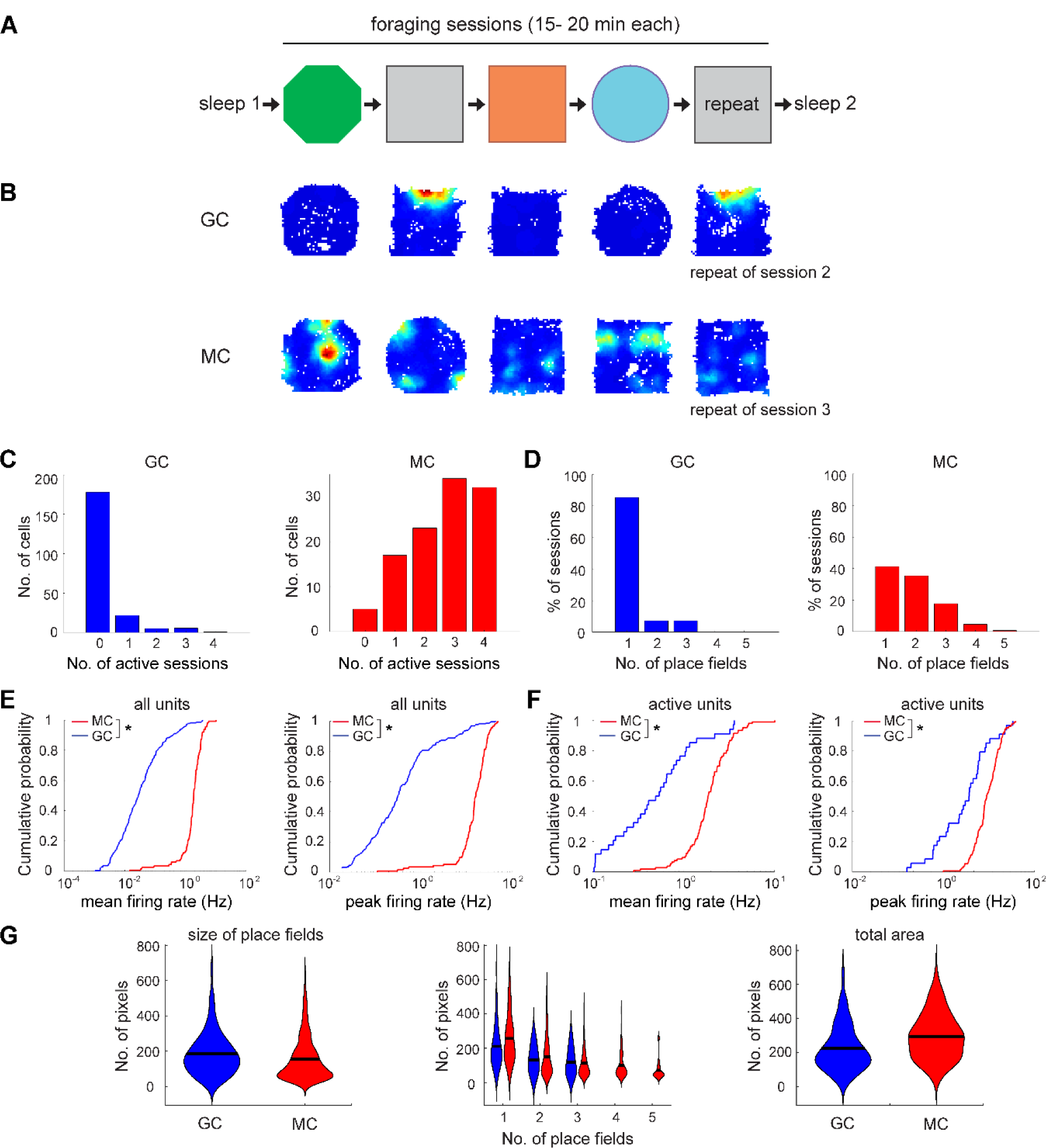
Experimental paradigm and general properties of classified granule cells and mossy cells. (A) Schematic of the experimental paradigms including sleep and behavior sessions in distinct environments. (B) Firing rate maps of a granule cell (GC) and a mossy cell (MC) that were recorded in four distinct environments and a repeat of an earlier session. Blue pixels in rate maps indicate areas with zero spikes, and red pixels reflect the cell’s peak firing rate across sessions. (C) The number of sessions with a place field for granule cells (left, blue) and mossy cells (right, red). (D) The number of place fields in each recording session. (E) Cumulative distribution functions (CDF) of mean firing rates and peak firing rates in the most active session for all cells (all units). (F) Same as E but for cells with a field in at least one environment (active units). (G) Left to right: the size of the individual place fields of granule cells and mossy cells (left), the size of the place fields plotted against the number of place fields (middle), and the total size of all place fields (right). See also **Figure S1** and **Table S1**

Since mossy cells and granule cells are in close anatomical proximity and a single mossy cell can be recorded from electrodes located 300 µm away from each other (Henze and Buzsaki, 2007), it is difficult to determine the identity of cells in the DG based solely on the electrode location. However, recent studies have distinguished granule cells and mossy cells based on electrophysiological properties (GoodSmith et al., 2017; GoodSmith et al., 2019; Jung et al., 2019; Senzai and Buzsaki, 2017). We used a random forests classifier to identify granule cells and mossy cells based on firing features obtained during the post-behavior baseline sleep session. We adapted a classification strategy from our previous study in rats (GoodSmith et al., 2017) and data from that study were used as training data for a new classifier with modifications to account for species differences between mice and rats, notably the reduced number of granule cells that could be recorded on a single tetrode in mice relative to rats. Classification features for the mouse-specific classifier included mean firing rate, burstiness, and channel slope (see Methods; Figure S1C). This classifier had an out-of-bag error rate (an estimate of the classifier’s generalization error) of 2.3% (Figure S1D), suggesting the classifier could reliably distinguish granule cells and mossy cells. Using this classifier, we classified 420 DG excitatory cells as 242 granule cells and 178 mossy cells.

The random forests classifier consists of a large number of decision tree classifiers generated using a subset of data and features. Each decision tree is given one “vote” and the classification with the most votes is the output of the ensemble classifier. The percentage of “votes” that the final classification receives can be viewed as a measure of the confidence in that classification (a cell that is classified as a mossy cell by 100% of the decision trees is less likely to reflect a misclassified granule cell than if only 51% of decision trees classified it as a mossy cell) (Figure S1E). In our current data set, for 77% (323/420) of cells, ≥ 85% of decision tree classifiers had the same output (212 granule cells and 111 mossy cells) (Figures S1E-G). To minimize the impact of any classification errors, we restricted our primary analyses to these most confidently classified cells. We repeated all analyses using all classified cells and did not find any significantly different results (Figure S2; Tables S2 and S3).

We first quantified the overall rates of activity and place field properties in granule cells and mossy cells during foraging sessions. Similar to prior reports (Danielson et al., 2017; GoodSmith et al., 2017; Kim et al., 2020; Senzai and Buzsaki, 2017), granule cells and mossy cells showed dramatically distinct firing characteristics (Figures 1B and S3). While only 16% (34/212) of granule cells had at least one session with a place field, 95% (106/111) of mossy cells had place fields in at least one environment (χ ^2^ = 187.31, p = 1.23 x 10^-42^), and 29% (32/111) of mossy cells had place fields in all sessions (Figure 1C). For sessions with at least one place field, the proportion of mossy cell sessions having multiple place fields was larger than that of granule cell sessions (MC: 172/293, 59%; GC: 8/54, 15%; χ ^2^ = 35.18, p = 3.01 x 10^-9^) (Figure 1D). The average number of place fields per session was higher for mossy cells than granule cells (GC: 1.20 ± 0.08; MC: 1.87 ± 0.07; rank-sum test z = -5.61, p = 2.04 x 10^-8^). Both mean firing rate and peak firing rate in the most active session of the day, irrespective of whether place fields were present, were higher for mossy cells than granule cells (mean firing rate: GC: 0.17 ± 0.03 Hz; MC: 2.12 ± 0.13 Hz; rank-sum test z = -13.83, p = 1.72 x 10^-43^; peak firing rate: GC: 2.04 ± 0.38 Hz; MC: 18.04 ± 0.90 Hz; rank sum test z = -13.38, p = 7.67 x 10^-41^) (Figure 1E). These differences remained when analysis was restricted to active cells that had at least one place field in at least one session throughout the recording day (mean firing rate: GC: 0.81 ± 0.16 Hz; MC: 2.19 ± 0.13 Hz; rank-sum test z = -6.47, p = 1.01 x 10^-10^; peak firing rate: GC: 10.59 ± 1.74 Hz; MC: 18.53 ± 0.87 Hz; rank-sum test z = -4.92, p = 8.54 x 10^-7^) (Figure 1F).

Individual place fields were larger on average for granule cells than for mossy cells (GC: 185.80 ± 13.48 pixels; MC: 156.09 ± 4.64 pixels; rank-sum test z = 2.84, p = 4.48 x 10^-3^), but the total area of all place fields in each environment was larger for mossy cells than for granule cells (GC: 224.63 ± 16.15 pixels; MC: 295.27 ± 7.05 pixels; rank-sum test z = -4.17, p = 3.01 x 10^-5^) (Figure 1G).

Taken together, these data show that mossy cells were more active and had more place fields than granule cells, and they frequently fired in multiple environments, whereas the overwhelming majority of active granule cells had a place field in only one environment. These results are consistent with previous reports (Danielson et al., 2016; Danielson et al., 2017; GoodSmith et al., 2017; GoodSmith et al., 2022; Senzai and Buzsaki, 2017) and demonstrate that we were able to reliably and accurately identify granule cells and mossy cells in DG recordings in our experiments.

### Global remapping by both granule cells and mossy cells in distinct environments

To estimate the extent to which neural activity patterns of each cell type varied among different environments, we first compared the mean firing rates of granule cells and mossy cells for each pair of foraging sessions from the same day, excluding the final repeat session. Only session pairs with a place field in at least one of the two sessions were used for this comparison. In plots of firing rates across each pair of sessions, most mossy cells had nonzero firing rates in each session whereas granule cells often exhibited a firing rate close to zero in one of the two sessions (Figure 2A).

**Figure 2.**
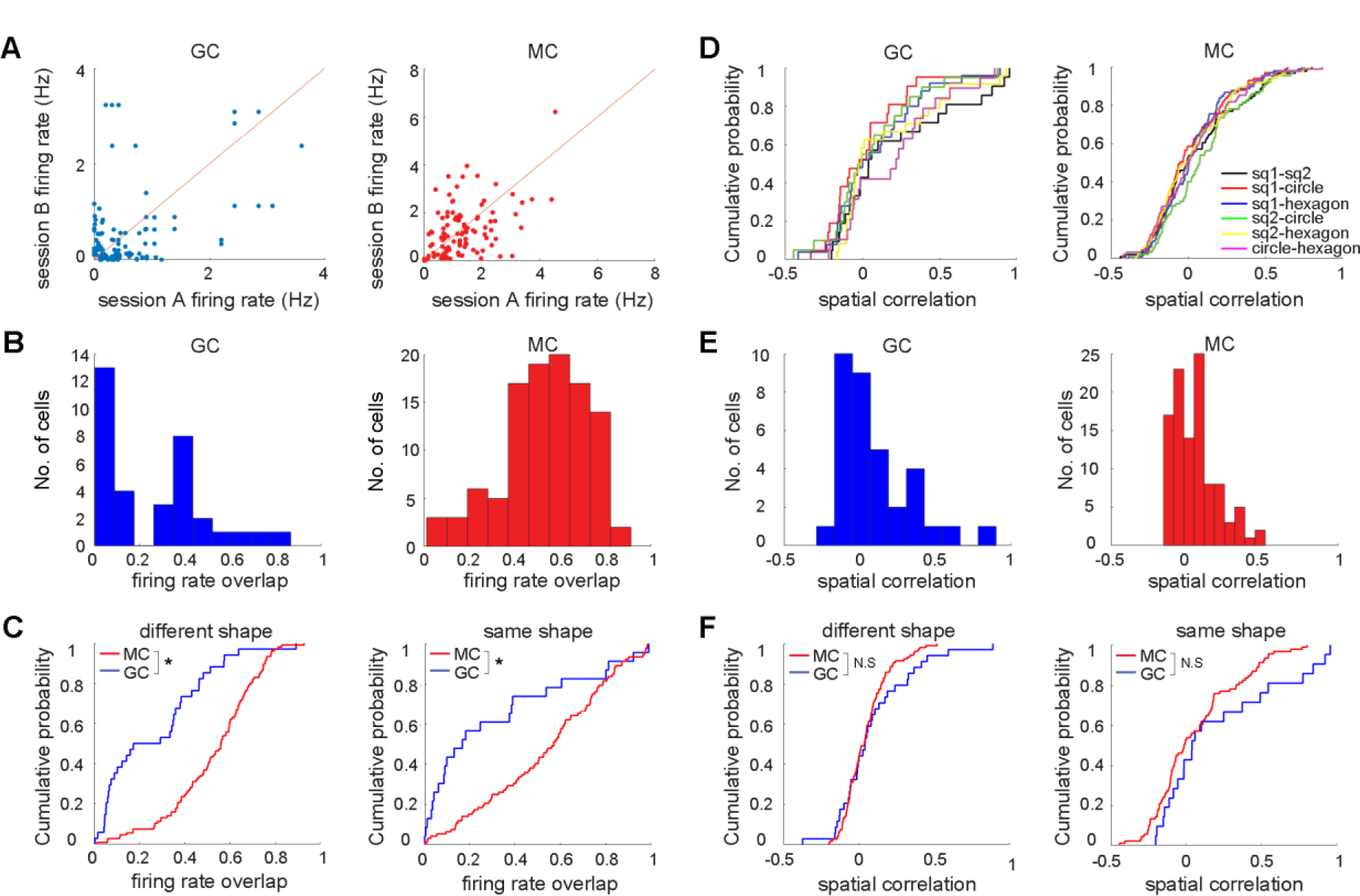
Rate overlap and spatial correlation comparison among four distinct sessions. (A) Comparison of mean firing rates from each session pair with a place field in at least one of the two sessions (4 environments, 3-6 pairs per cell). (B) Histograms of firing rate overlap for granule cells (left) and mossy cells (right). (C) Cumulative distribution functions (CDF) of the firing rate overlap for different shape (left) and same shape conditions (right) (granule cells (blue) and mossy cells (red)). (D) CDF of the spatial correlations for granule cells (left) and mossy cells (right) from each pair of sessions. (GC: sq1-sq2: 0.21 ± 0.08, sq1-circle: 0.03 ± 0.06, sq1-octagon: 0.09 ± 0.06, sq2-circle: 0.06 ± 0.07, sq2-octagon: 0.07 ± 0.20, circle-octagon: 0.20 ± 0.07. Kruskal-Wallis: z =5.32, p = 0.38; MC: sq1-sq2: 0.07 ± 0.03, sq1-circle: 0.02 ± 0.02, sq1-octagon: 0.04 ± 0.02, sq2-circle: 0.11 ± 0.03, sq2-octagon: 0.04 ± 0.03, circle-octagon: 0.04 ± 0.03. Kruskal-Wallis: z =6.70, p = 0.24) (E) Histogram of the spatial correlations for granule cells (left) and mossy cells (right). (F) CDF of the spatial correlation for different shape (left) and same shape conditions (right) for granule cells (blue) and mossy cells (red). See also Figures S2 and S3 and **Table S1**.

To quantify the similarity in firing rates, rate overlap values were calculated by dividing the smaller mean firing rate by the larger mean firing rate from all session pairs with at least one field in one environment (Leutgeb et al., 2007) (Figure 2B). A rate overlap value of 1 occurs when the firing rate is the same in both environments, while a value closer to zero indicates that a cell fired predominately in one of the two environments. Granule cells typically had place fields in a single environment and mossy cells frequently had place fields in both environments, resulting in significantly higher rate overlap values for mossy cells than granule cells (GC: 0.26 ± 0.04; MC: 0.53 ± 0.02; rank-sum test z = -5.69, p = 1.29 x 10^-8^) (Figure 2B). To examine how these cell types respond to different degrees of environmental changes, we used two different manipulations: manipulation of environment shape, color and floor texture (different shape condition), or manipulation of color and floor texture only, using the same square geometry (same shape condition) (see Methods). We found that the rate overlap values for granule cells were significantly lower than those obtained from mossy cells for both different shape and same shape conditions (different shape: GC: 0.27 ± 0.04; MC: 0.53 ± 0.02; rank-sum test z = -5.54, p = 3.07 x 10^-8^; same shape: GC: 0.30 ± 0.07; MC: 0.54 ± 0.03; rank-sum test z = -3.25, p = 1.13 x 10^-3^) (Figure 2C). These data are consistent with the general properties of each cell type, with sparsely firing granule cells showing the largest changes in firing rate due to the increased likelihood of having a place field in only one environment in any pair of sessions.

We next examined how different environments altered the spatial firing patterns of granule cells and mossy cells. We calculated the spatial correlation values for all 6 combinations of the four distinct environments and found that each combination of session pairs with a field in at least one session had similar spatial correlation values for both granule cells and mossy cells (Figure 2D). Among the four distinct environments, both granule cells and mossy cells exhibited strong remapping due to low spatial correlation and there was no significant difference in the degree of remapping between the two cell types (spatial correlation: GC: 0.11 ± 0.04; MC: 0.05 ± 0.01; rank-sum test z = 0.62, p = 0.54) (Figure 2E) or between male and female mice for either cell type (Table S1). We also found that spatial correlation values for granule cells were similar to those obtained from mossy cells for both different shape and same shape conditions (different shape: GC: 0.09 ± 0.04; MC: 0.05 ± 0.01; rank-sum test z = 0.46, p = 0.65; same shape: GC: 0.21 ± 0.08; MC: 0.07 ± 0.03; rank-sum test z = 1.42, p = 0.15) (Figure 2F). Taken together, our data indicate that there is no substantial difference in the degree of remapping between granule cells and mossy cells as measured by spatial correlation, but that there is a significant difference between the populations in terms of firing rate changes across sessions.

### Reproducibility and remapping in granule cells and mossy cells

To determine the extent to which firing patterns were reinstated upon re-exposure to the same environment, the final session of the day was a repeated exposure to one of the four previous environments. We compared both the rate overlap and spatial correlation values in all pairs of sessions with a field in at least one session to measure remapping in session pairs with spatially modulated firing. Both granule cells and mossy cells showed higher rate overlap values for the repeat session pairs than the different environment pairs, suggesting a certain degree of stability of firing upon re-exposure to the same environment (GCs: repeat: 0.51 ± 0.07; different environment: 0.21 ± 0.03; rank-sum test z = 3.39, p = 6.88 x 10^-4^; MCs: repeat: 0.65 ± 0.03; different environment: 0.55 ± 0.02; rank-sum test z = 3.47, p = 5.29 x 10^-4^) (Figure 3A).

**Figure 3.**
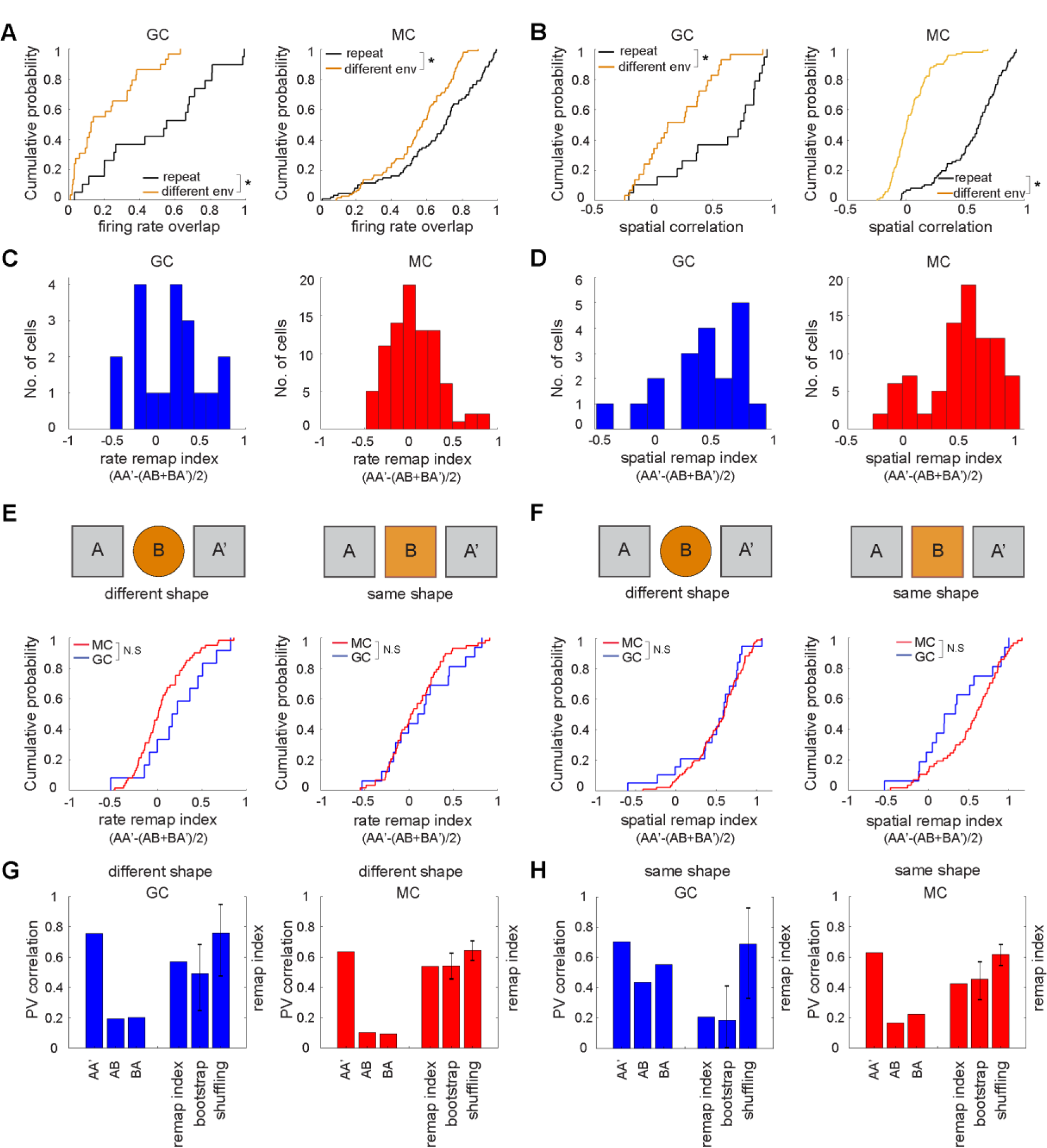
Rate overlap and spatial correlation comparison with a repeat session. (A) CDF of the rate overlap for granule cells (left) and mossy cells (right) (repeated session: black, different environment: orange). (B) CDF of the spatial correlation for granule cells (left) and mossy cells (right) (repeated session: black, different environment: orange). (A-B) Only session pairs in which the cell had a place field in at least one of the two environments were included for analysis. (C) Histograms of the rate remapping index for granule cells (left) and mossy cells (right). Remapping index. (AA’-(AB+BA’)/2). (D) Histograms of the spatial remapping index for granule cells (left) and mossy cells (right). Remapping index. (AA’-(AB+BA’)/2). (E) CDF of the rate remapping index for different shape (left) and same shape conditions (right) (granule cells (blue) and mossy cells (red)). (F) CDF of the spatial remapping index for different shape (left) and same shape conditions (right) (granule cells (blue) and mossy cells (red)). (G-H) Population vector (PV) correlation for the (G) different shape and (H) same shape environment. PV correlation was calculated for AA’, AB, BA’ and the remapping index was calculated by subtracting the average of AB and BA’ values from the AA’ value. Shuffled distributions were generated by shuffling cell ID in session B 1000 times and bootstrapping was performed 1000 times by using the whole population of granule cells and mossy cells. Error bars represent lower and upper bounds of 95% two-tailed confidence interval. (C-H) Remapping index was calculated by using subpopulations with place fields in either A or A’. See also **Figures S2, S3 and S4**.

Similarly, for spatial correlation values, both granule cells and mossy cells exhibited higher correlation values for the repeat session pairs than the different environment pairs, indicating that both cell types maintained a stable spatial representation during the second exposure to an environment (GCs: repeat: 0.56 ± 0.09; different environment: 0.20 ± 0.06; rank-sum test z = 3.01, p = 2.57 x 10^-3^; MCs: repeat: 0.56 ± 0.03; different environment: 0.04 ± 0.02; rank-sum test z = 10.04, p = 1.05 x 10^-23^) (Figure 3B).

Next, we adapted a measure from a previous study (Senzai and Buzsaki, 2017) to calculate a remapping index across a trio of sessions: the first environment (session A), a second environment (session B), and a repeat session of the initial environment (session A’). The key modification we made was the criterion used to select cells for analysis. We required that the cell must have a field in either A or A’, whereas the previous study only required that a cell have a field in at least one of the three sessions; that is, under our criterion, a cell that only had a field in session B would not be included, whereas it was included in the analysis of Senzai and Buzsaki. We made this modification because the previous criterion allowed for the inclusion of a correlation between two sessions in which the cell hardly fired (A and A’), which, in principle, might have added random noise to the data by correlating two weak rate maps with random spatial firing, thereby artificially reducing the overlap index for the A vs. A’ comparison based on that noise.

We first applied the index to rate remapping, which was defined as the difference in rate overlap values between the repeated session (AA’) and the average of the session pair that included the first exposure to environment A (AB) and the session pair with the second exposure to environment A (BA’) (Senzai and Buzsaki, 2017). Using this rate remapping index with our modified criterion, we found no significant differences in the magnitude of rate remapping between the two cell types (GC: 0.14 ± 0.09; MC: 0.06 ± 0.03; rank-sum test z = 0.95 p = 0.34) (Figure 3C). To investigate the effect of changing the geometry of the environment, we compared the rate remapping index from the different shape and same shape conditions for each cell type. We found that the rate remapping index values from granule cells were not significantly different from mossy cells for either the different shape or same shape pairs (different shape GC: 0.21 ± 0.11; MC: 0.04 ± 0.04; rank-sum test z = 1.64, p = 0.10; same shape GC: 0.15 ± 0.10; MC: 0.08 ± 0.04; rank-sum test z = 0.51, p = 0.61) (Figure 3E).

We next calculated the spatial remapping index for these same cells with our modified inclusion criterion. The spatial remapping index had the same form as the rate remapping index, but used spatial correlation values instead of rate overlap values. The spatial remapping index values of granule cells were not significantly different from those of mossy cells (GC: 0.44 ± 0.09; MC: 0.52 ± 0.03; rank-sum test z = -0.88 p = 0.38) (Figure 3D). We also found no significant difference in the spatial remapping index between the two cell types for both different shape and same shape conditions (different shape GC: 0.46 ± 0.09; MC: 0.53 ± 0.03; rank-sum test z = -0.45, p = 0.66; same shape GC: 0.32 ± 0.11; MC: 0.53 ± 0.05; rank-sum test z = -1.63, p = 0.10) (Figure 3F). Although not significant, granule cells exhibited a trend toward lower remapping index scores than mossy cells in the same shape condition, suggesting that granule cell remapping may be sensitive to the degree of change in the environment.

Given that we observed similar levels of remapping in both granule cells and mossy cells using our modified inclusion criterion, we then recalculated the remapping indices using the same criterion as the previous study (Senzai and Buzsaki, 2017) and included all cells with a place field in A, B, or A’. There was no significant difference between the two cell types in the rate remapping index, but the spatial remapping index was lower for granule cells than for mossy cells (Figure S4 and Table S4). This difference between granule cells and mossy cells was similar to what was previously reported (Senzai and Buzsaki, 2017) and may be partly driven in our data set by random noise introduced when quantifying correlations between sessions with very weak firing, which occurs more often in granule cells than mossy cells (Figure S4).

To analyze remapping properties at the population level, we quantified the population vector (PV) correlations of granule cells and mossy cells across a trio of sessions (A-B-A’) (Figures 3G and 3H, left) (Leutgeb et al., 2007; Senzai and Buzsaki, 2017). We also calculated the remap index using the PV correlation values (Figures 3G and 3H, right). While the remap index value for granule cells was similar to that of mossy cells for the different shape condition, the remap index value of granule cells was lower than that of mossy cells for same shape condition (different shape GC: 0.57; MC: 0.54; same shape GC: 0.21; MC: 0.43) (Figures 3G and 3H). To estimate the distribution for each population, we performed bootstrapping by sampling all of the values for each cell type. For the different shape condition, the lower bound of the 95% confidence interval of the bootstrapped values for both mossy cells and granule cells was a positive value, indicating that there was remapping in both populations (GC: remap index: 0.49, lower bound: 0.25, upper bound: 0.68; MC: remap index: 0.54, lower bound: 0.46, upper bound: 0.63) (Figure 3G).

For the same shape condition, the 95% confidence interval of the bootstrap distribution for mossy cells encompassed only positive values, whereas the lower bound of the 95% confidence interval for granule cells was -0.004 (29/1000 bootstrapped samples were < 0), suggesting that the evidence for remapping of granule cells was marginal in this analysis (GC: remap index: 0.19, lower bound: 0, upper bound: 0.41; MC: remap index: 0.45, lower bound: 0.32, upper bound: 0.57) (Figure 3H).

Finally, to investigate whether any differences in the degree of remapping may be related to inherent biases introduced by the unique firing properties of granule cells and mossy cells, we shuffled the cell identity in session B to simulate complete, global remapping and calculated the remap index values from these shuffled populations. The remap index values of the shuffled distributions of granule cells were not lower than those of mossy cells, indicating that the lower degree of remapping in granule cells in the same shape condition was not derived from intrinsic properties of granule cells (different shape: GC: remap index: 0.76, lower bound: 0.48, upper bound: 0.95; MC: remap index: 0.65, lower bound: 0.58, upper bound: 0.71; same shape: GC: remap index: 0.69, lower bound: 0.33, upper bound: 0.93; MC: remap index: 0.62, lower bound: 0.55, upper bound: 0.68) (Figures 3G and 3H). The actual remap index values for both cell types were lower than the 95% confidence interval for their respective shuffled distributions, except for granule cells in the different shape condition, indicating less than complete remapping in most conditions (Figures 3G and 3H). Our data suggest that granule cells and mossy cells remapped to a similar degree in the different shape condition, while granule cells remapped less when the two environments shared the same shape.

### Granule cell and mossy cell populations remap to a similar extent, but with different properties

To more fully represent how the populations of granule cells and mossy cells modify their activity across sessions, we next quantified the proportion of session pairs with place fields in either one session or both sessions. In total, there were 133 session pairs in which granule cells had a place field in at least one session. Among these, 22% (29 out of 133) of session pairs had a place field in both sessions, indicating that the majority of granule cell session pairs with place fields (78%) had fields in only one session. This result suggests that remapping in granule cells is mediated largely by the recruitment of distinct populations of cells with place fields to represent each environment. In contrast, among mossy cell session pairs having place fields, more than half of the session pairs (56%, 317 out of 562) had fields in both sessions.

We next quantified remapping using a correlation value threshold to set a cutoff for what was considered remapping due to spatial decorrelation (Figures 4A and 4B). Given that the average spatial correlation value for repeat sessions was 0.56 for both cell types, we used one standard deviation away from this mean value as a threshold to define remapping (GC threshold: 0.18; MC threshold: 0.29). Only session pairs having a place field in both sessions were included for this analysis. Any session pair with a spatial correlation value below this threshold was categorized as ‘spatially decorrelated’, whereas session pairs with correlation values higher than this cut-off were defined as ‘spatially correlated’. We found that 59% of granule cell session pairs (17 out of 29) and 83% of mossy cell session pairs (264 out of 317) with fields in both environments showed lower spatial correlation values than the cut-off and were categorized as ‘spatially decorrelated’ (Figure 4C).

**Figure 4.**
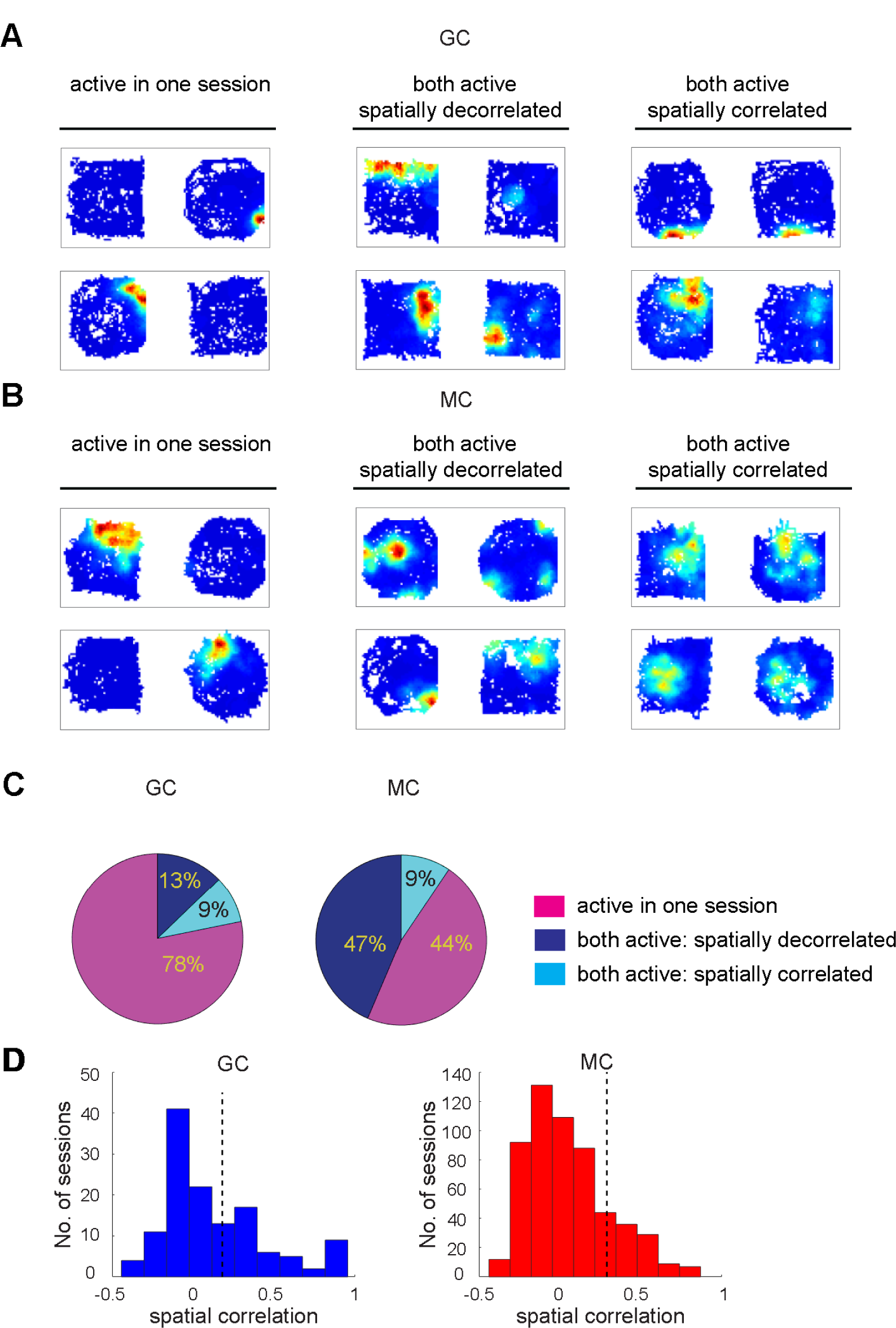
Different patterns of global remapping. (A) Firing rate map examples of granule cell session pairs with place fields in one session (left), with place fields in both sessions but with low spatial correlation (middle), and with place fields in both sessions with stable place fields (right). (B) Same as A but for mossy cells. (C) Proportion of session pairs with place fields in one session only (active in one session) or with place fields in both sessions that either meet the correlation threshold (spatially correlated) or fail to meet the correlation threshold (spatially decorrelated). (D) Histogram of the spatial correlations for granule cells (left) and mossy cells (right) with a black line indicating the threshold for spatially correlated sessions (GC: 0.18, MC: 0.29).

Both granule cells and mossy cells had a similar percentage of spatially correlated sessions (9%, Figure 4C), but we found that several granule cell session pairs had very high spatial correlation values (6 session pairs > 0.8), suggesting that a small subset of the granule cell population exhibited highly stable spatial firing across different environments (Figure 4D). This finding replicates observations from a previous study showing stability in a small percentage of granule cells (Hainmueller and Bartos, 2018). In total, 91% of the session pairs having at least one place field exhibited some form of remapping by both granule cells (121 out of 133) and mossy cells (509 out of 562), either due to a low correlation between place fields in the two environments (spatial decorrelation) or by having a place field(s) in one environment but not the other (active population remapping) (Figure 4C). This result indicates that the two cell types remapped to a similar extent. Interestingly, although a similar proportion of granule cells and mossy cells exhibited remapping, the proportions of remapping through turnover in the active population vs. spatial decorrelation of place fields across environments were different between granule cells and mossy cells (χ ^2^ = 56.58, p = 5.39 x 10^-14^): a higher proportion of granule cell than mossy cell session pairs exhibited remapping due to turnover in the active population (GC: 104/121 (86%); MC: 245/509 (48%)) and, conversely, a higher proportion of mossy cell than granule cell session pairs showed remapping by spatial decorrelation (MC: 264/509 (52%); GC: 17/121 (14%)). Taken together, these data suggest similar levels of remapping in granule cells and mossy cells but differences in the population-level mechanisms of remapping (GoodSmith et al., 2017).

## DISCUSSION

Remapping of hippocampal place cell activity reflects a neural correlate of context specificity that may support the formation of episodic memories. Within the hippocampus, the DG has long been hypothesized to be critical for encoding context specificity through pattern separation.

However, recent studies have called into question the capacity of granule cells to exhibit strong global remapping in response to changes in the environment in virtual reality (Hainmueller and Bartos, 2018) or in environments that share the same physical location (Senzai and Buzsaki, 2017). Here we resolved controversies over remapping in the DG and demonstrate robust remapping in granule cells. We recorded granule cells and mossy cells as mice freely explored four different environments located in the same location and were re-exposed to one environment each day. Using a random forests classifier to discriminate between cell types, we demonstrated differences in the spatial firing properties of these populations, as granule cells typically exhibited only one, if any, place field(s) in one environment, whereas mossy cells fired much more promiscuously, often with multiple place fields in multiple environments (GoodSmith et al., 2017; GoodSmith et al., 2022; Senzai and Buzsaki, 2017). These differences influenced how each cell type contributed to global remapping and required analyses that reflected the underlying properties of spatially modulated firing in each population. By including multiple measures of remapping, we were able to account for the sparse firing and high degree of turnover in the population of granule cells that exhibited spatially modulated activity in each environment. These results support a role for both excitatory cell types of the DG in pattern separation and the flexible encoding of spatial information needed to support context-specific memory formation.

### Robust remapping in granule cells

Recent studies have reported widely varying levels of global remapping in identified granule cells (Allegra et al., 2020; GoodSmith et al., 2017; Hainmueller and Bartos, 2018; Senzai and Buzsaki, 2017). Previous studies of freely moving animals differed in the degree to which the contextual features were altered in each environment, which could at least partially account for the discrepancy in magnitude of remapping. One study used different shaped arenas in physically distinct rooms and global remapping was observed in both granule cells and mossy cells

(GoodSmith et al., 2017); another used two identically shaped arenas in the same location with differences in the local cues on the walls and floor and reported lower levels of remapping in granule cells than mossy cells (Senzai and Buzsaki, 2017). To determine whether changing the geometry of the enclosure influenced remapping, we used four distinct arenas, two of which had a similar square shape. Both mossy cells and granule cells showed robust global remapping when the shape of the environment was modified in addition to the other features. In the same shape environments, our data showed a trend toward lower remapping in granule cells than mossy cells across several measures including the spatial remapping index and population vector correlation. This suggests that granule cells are sensitive to the degree of change across environments and can remap less than mossy cells when prominent features of the environment overlap. Senzai and Buzsaki (2017) changed visual wall cues and floor texture in an otherwise identically shaped and colored apparatus to create two contexts; they reported a modest level of remapping in granule cells compared to more robust remapping in mossy cells. In our same-shape condition, we showed evidence for remapping of granule cells in the distribution of rate remap and spatial remap indices (Figures 3E and 3F). In addition, the PV remap index was 0.19, with only 29 out 1000 bootstrapped samples producing a remap index < 0 (Figure 3H); thus, the 2-tailed 95% confidence interval barely included the null value of 0. Given the strong a priori prediction that the remap index would be non-negative (i.e., it is very unlikely that the rate maps would be more correlated between different environments, A and B, than between repeated exposures to the same environment, A and A’), a reasonable argument can be made that a one-tailed test is appropriate, in which case the null value of 0 falls outside the 95% confidence interval of the bootstrap. Moreover, the rate remap index and spatial remap index were not significantly different between granule cells and mossy cells (Figure 3E and 3F). Given the small proportion of active granule cells with place fields in each study, we suggest that there is a small but detectable amount of remapping of granule cells in same-shape conditions in both the present study and the study of Senzai & Buzsaki (2017), and that the difference in magnitude between the studies may be a result of sampling error. Furthermore, the incremental increase in granule cell remapping that accompanies incremental changes in the environment (Knierim and Neunuebel, 2016) may induce the robust global remapping of mossy cells in both same- and different-shape conditions, as the mossy cells receive small remapping signals from granule cells in the same shape environment and amplify these “seed” signals as part of a larger DG pattern separation circuit with backprojections from proximal CA3 (GoodSmith et al., 2019; Hunsaker et al., 2008; Lee et al., 2015; Myers and Scharfman, 2009; 2011; Scharfman, 2007; Senzai and Buzsaki, 2017).

Another study that reported low levels of remapping in granule cells greatly underestimated the extent of that remapping by emphasizing the small proportion of granule cells that had place fields in both environments. (Hainmueller and Bartos, 2018). This study reported that ∼10% of the granule cell population had place field activity that was highly correlated across two different environments and sustained over several days (Hainmueller and Bartos, 2018).

Based on this subset of cells, the authors argued that granule cells are more stable than CA1 pyramidal cells and did not remap across environments. However, this interpretation failed to account for the robust remapping due to the high turnover in the population of granule cells with place fields in each environment. In their data set, ∼90% of granule cells with place fields were active in only one of the two environments, demonstrating strong remapping by nonoverlapping active populations, which is highly consistent with our results and constitutes very strong, global remapping at the population level. This interpretation was strongly supported by Allegra et al. (2020) under conditions similar to the Hainmueller and Bartos (2018) study.

### Reproducibility of spatial maps in DG cells

While remapping may contribute to discrimination among different environments, the reproducibility of spatial maps during repeated exposures to a single environment can support the recognition of familiar contexts, which is also critical for the formation of episodic and associative memory (Dupret et al., 2010; O’Keefe and Nadel, 1978). Compared to place cells in rats, place cells in mice have been shown to be unstable in specific tasks (Dong et al., 2021; Gonzalez et al., 2019; Hussaini et al., 2011; Kentros et al., 2004; Levy et al., 2021; Ziv et al., 2013) and the stability of spatially-modulated activity also varies among hippocampal subregions (Dong et al., 2021; Hainmueller and Bartos, 2018; Mankin et al., 2015; Schoenfeld et al., 2021). To measure stability of DG spatial representations in mice in our study, we evaluated activity patterns in both granule cells and mossy cells upon re-exposure to one environment at the end of the recording day. Spatial correlation values were significantly higher between repeated exposures to a single environment than between different environments for both granule cells and mossy cells, suggesting that spatial representations in the DG can be at least partially reinstated, even following exposure to intervening environments that induced remapping.

Interestingly, we found that a small subset of granule cells (3 out of 34, ∼10%) showed a very high spatial correlation value (>0.8) across multiple pairs of sessions in different environments. This result is consistent with a previous finding that ∼10% of granule cells with place fields maintain a similar spatial firing pattern across environments (Hainmueller and Bartos, 2018). It is not clear whether these cells represent a distinct subpopulation of granule cells. Given that the stable cells represent only ∼10% of the granule cell population, it seems unlikely that these are developmentally born granule cells, which comprise the majority of the granule cell population. One possibility is that these cells are adult-born immature granule cells, which have been proposed to be more active during a transient period of maturation (Ming and Song, 2011). Increased intrinsic excitability in immature granule cells could lead to more permissive gating of dendritic inputs and a higher probability of firing in response to inputs (Diamantaki et al., 2016; Krueppel et al., 2011; Schmidt-Hieber et al., 2007; Zhang et al., 2020). However, most manipulations of adult-born neurons have suggested immature granule cells play a prominent role in mediating behavioral context discrimination (Besnard and Sahay, 2021; Nakashiba et al., 2012), which may be at odds with firing properties that are invariant to contextual changes. Further, imaging studies have suggested that immature granule cells exhibit less spatial selectivity than mature granule cells overall, but can still discriminate among environments based on changes in spatial tuning (Danielson et al., 2016). Future studies are needed to determine whether there are other characteristic properties of these cells and whether they represent a subpopulation of all granule cells or granule cells of a specific developmental age. Together, these results suggest that a small subset of granule cells are more likely to fire in multiple environments with less context specificity, whereas the majority of spatially tuned granule cells remap by alternating activation states among environments through the presence or absence of a place field.

## Conclusion

Global remapping emerges from unique ensembles of cells that are active in each environment, as well as the independent spatial representations of cells that have place fields in more than one environment (Bostock et al., 1991; Knierim, 2003; Kubie and Muller, 1991). Our data show that mossy cells are more likely to remap by *reorganization of place fields* across environments and granule cells are more likely to remap by *reorganization of active populations* across environments. As predicted by theory (Yassa & Stark 2011; Knierim & Neunuebel 2016), the amount of remapping of granule cells can vary depending on the amount of change to its inputs (not all of which are under direct experimental control or measurement) before granule cells create completely orthogonal representations when the change in inputs is sufficiently large.

This variability, combined with differences in analysis methods, can explain the different levels of granule cell remapping reported by different experiments. Overall, the literature strongly supports a role for both granule cells and mossy cells in flexible spatial encoding through distinct patterns of firing across different environments and provides strong support for the classic theory that a major—although probably not exclusive—function of the dentate gyrus (GoodSmith et al., 2019; Lee et al., 2020; Lee et al., 2015; Lee et al., 2022) is to perform pattern separation (Hunsaker et al., 2008; Jinde et al., 2012; Marr, 1969; McNaughton and Nadel, 1990; Nakashiba et al., 2012; Rolls and Kesner, 2006; Yassa and Stark, 2011).

## ACKNOWLEDGMENTS

We thank B. Temsamrit, E. LaNoce, A. Garcia, and A. Angelucci for technical support and lab coordination and S. Mysore for advice on the classification method. This work was supported by grants from the National Institutes of Health (R01NS039456 to J.J.K., R35NS116843 to H.S., and R35NS097370 to G-l.M.) and Dr. Miriam and Sheldon G. Adelson Medical Research Foundation (to G.-l.M.).

## AUTHOR CONTRIBUTIONS

S.H.K., K.M.C., H.S. and J.J.K. designed the experiments; S.H.K., D.G. and S.J.T. collected the data; S.H.K., D.G., F.M. and K.M.C. analyzed the data; K.M.C., G-l.M., H.S., and J.J.K. provided supervision and funding; S.H.K. and K.M.C. wrote the initial draft of the manuscript, and S.H.K, K.M.C., and J.J.K. wrote the final version; and all authors provided comments and feedback on the manuscript.

## STAR METHODS

### Key resources table

| **REAGENT or RESOURCE** | **SOURCE** | **IDENTIFIER** |
| --- | --- | --- |
| VECTASHEILD<br>Mounting medium<br>with DAPI | Vector<br>Laboratories | Cat#H-1500 RRID AB_236788 |
| Matlab | Mathworks | https://www.mathworks.com/products/matlab |
| Digital Lynx SX data<br>acquisition system | Neuralynx | Neuralynx Neuralynx.com |
| Cheetah acquisition<br>software | Neuralynx | Neuralynx Neuralynx.com |
| Violin plot | Holger Hoffmann | https://www.mathworks.com/matlabcentral/fileexchange/45134-violin-plot |

### Resource availability

*Lead contact:* Requests for further information, data, and other resources can be directed to the lead contact, James J. Knierim.

*Materials availability:* This study did not generate new materials.

*Data and code availability:* All data and analysis code will be available on GitHub (GitHub: https://github.com/skim1011/KimGoodSmith_DG_remap).

### Methods details

#### Subjects and surgery

Neural activity was recorded in dentate gyrus (DG) from 35 C57BL/6 mice (23 male, 12 female, 19-32 g, 10-35 weeks old, Tbr2-CreERT2::ChR2 transgenic line). All experiments were performed in accordance with National Institutes of Health guidelines and approved by the Institutional Animal Care and Use Committees at Johns Hopkins University and the University of Pennsylvania. During surgery, mice were anesthetized with a mixture of ketamine/xylazine/acepromazine (70-100, 5-12, and 1-3 mg/kg, respectively) or isoflurane (0.5%-2.5% in O2). Custom-built recording drives with 7 tetrodes and an optic fiber were implanted centered at 2.0 mm posterior to bregma and 1.3 mm lateral to the midline (Kloosterman et al., 2009). A 200 µm diameter optic fiber was inserted above the granule cell layer (DV 1- 1.5) and blue light pulses (≤ 5ms per pulse) were delivered after the post-behavior sleep session for optical tagging of adult-born neurons for another experiment. After surgery, mice were individually housed on a 12-h light/ dark schedule with *ad libitum* access to water.

#### Training and behavior

After recovery from surgery, mice were trained to forage for small chocolate sprinkles (Sprinkle King) in 4 distinct, enclosed environments, including a circular platform, an octagon platform, and two square platforms with different colors and textures (squares: 60 cm width, 30 cm height; octagon 64 cm total width (four edges each at 22 cm and 33 cm length), 17 cm height; circle: 69 cm width, 8 cm height). Each environment had distinct cues, different colors, patterns and textures on the walls and floors (square 1: black paper on the floor and circle cue on the red wall. square 2: brown artificial leather floor and square cue on the brown wall. circle: blue pattern paper on the floor and circle cue on the metal wall. octagon: black mesh on the floor and X shape cue on the white wall.)

#### Electrophysiological recordings

Tetrodes were constructed from 12 µm or 17 µm nichrome wire (California Fine Wire, Grover Beach, CA). Nichrome wires were gold-plated to reduce wire impedance to the range between 150-400 kΩ. During recording, a custom-designed hyperdrive was connected to a Digital Cheetah Data Acquisition System (Neuralynx, Bozeman, MT). The signal was amplified 1000- 5000 times and filtered between 600 Hz and 6 kHz (for single units) or 1 Hz and 475 Hz (for local field potentials). Spike waveforms above a threshold (40-70 µV) were sampled at 32 kHz for 1 millisecond.

#### Histological procedures

The mice were anesthetized with a mixture of ketamine/xylazine/acepromazine (70-100, 5-12, and 1-3 mg/kg, respectively) and were perfused with phosphate-buffered saline, followed by 4% paraformaldehyde (PFA) in saline. The brains were cut in coronal sections (40-50 µm) with a microtome, mounted on glass slides, and stained with DAPI (Vector Laboratories). Sections were viewed and imaged using a confocal microscope (Zeiss) to identify tetrode tracks and confirm location in the DG.

#### Unit isolation

Single units were isolated with custom-written, manual cluster-cutting software (Winclust, J. Knierim) using multiple waveform characteristics (e.g., spike amplitude peak and energy under the waveform). The isolation quality of single units was subjectively rated with a scale from 1 (very good) to 5 (poor). Unit isolation rating was made independently of the behavioral firing correlates of the cells. The cells rated as marginal or poor (4 and 5) were excluded from further analysis.

#### Rate maps and place fields

The position of the mouse during the foraging session was detected by video tracking of red and green LEDs on the headstage. Firing rate maps were generated by dividing the number of spikes of a single cell in each bin (64 x 48) by the total time the mouse spent in that bin (pixel size ∼1.3 cm^2^). Rate maps were speed filtered (> 2 cm/s) and smoothed with an adaptive binning method (Skaggs et al. 1996). The mean firing rate, peak firing rate, and the spatial information score (Skaggs et al. 1996) were calculated from these rate maps. The significance of the spatial information score was determined by a shuffling procedure, in which the spike train and position train were shifted relative to each other by a random amount (minimum of 30 seconds), and the spatial information score was recalculated based on a regenerated rate map. This procedure was performed 100 times, and the spatial information score was considered significant at p < 0.01 if the observed information score was higher than all shuffles. Place cells were defined as cells with mean firing rate ≥ 0.1 Hz, spatial information score > 0.5 bits/spikes, and p-value for spatial information < 0.01. A place field was identified as a region with ≥ 30 contiguous pixels that exceeded 20% of the peak firing rate. Putative high-rate interneurons with a mean firing rate more than 10 Hz during the sleep session were excluded from analysis. There was an additional population of cells with a mean firing rate between 2 and 10 Hz but no spatial firing in any given environment. These cells may comprise a different population of putative interneurons in the hilar region (GoodSmith et al., 2017; Houser, 2007) and were also excluded from analysis.

#### Rate overlap and spatial correlation analysis

Firing rate overlap scores between two sessions were calculated by dividing the mean firing rate in the session having the lower firing rate by the mean firing rate in the session having the higher firing rate. Spatial correlation was calculated using Pearson’s correlation between the firing rates of each bin from two rate maps. Unvisited pixels in either session were excluded from the calculation. Cells with place fields in at least one environment were considered for rate overlap and spatial correlation analysis, and session pairs that had a place field in at least one of the two sessions being compared were included.

Three sessions in A-B-A’ format were used to determine the remapping index (A: initial environment, B: different environment, A’: a repeat session of the initial environment). The remapping index was calculated by subtracting the average of AB and BA’ values from the AA’ value (rate overlap values for the rate remap index, spatial correlation values for the spatial remap index, and PV values for the PV remap index). Cells having a place field in either A or A’ were included for the remapping index analyses.

Only units having a place field in either A or A’ were included for population vector analysis. All rate maps were stacked into a three-dimensional matrix with x and y axes containing firing rate information. The remap index based on population vector (PV) correlations was obtained by calculating the difference between the AA’ PV correlation and the mean of AB and BA’ PV correlations. The remap index was also calculated for a bootstrapped distribution of 1000 samples with replacement for each cell type and by shuffling cell identity in session B 1000 times.

#### Cell type classification

Cells recorded in the DG were classified as granule cells or mossy cells based on their firing in the post-behavior baseline/sleep session. To classify cells into putative cell types, a random forests classifier was created (Brieman, 2001; Liaw and Wiener, 2002). Random forests classifiers are generated by creating a large number of decision tree classifiers, producing an ensemble classifier that is less prone to overfitting than other supervised learning methods. Each decision tree is created using a bootstrapped sample of training data and a random subsample of classification features. The overall output of the classifier is defined as the mode of individual classifier outputs (each decision tree is given one “vote” and the output with the most votes is the final classifier output).

As training data for the classifier, we used previously published recordings from the DG in the rat (GoodSmith et al., 2017). In that study, a total of 130 cells were recorded on tetrodes located clearly in the DG (excluding any tetrode that was closer to the CA3c pyramidal cell layer than the granule cell layer). For our training data, we considered all cells from this dataset that had place fields in zero or one environment as putative granule cells. We considered any cells with a place field in at least 3 environments to be putative mossy cells. This putative cell type assignment was based on the observed firing properties of granule cells and mossy cells in rats (Diamantaki et al., 2016; GoodSmith et al., 2017; GoodSmith et al., 2019; Leutgeb et al., 2007; Neunuebel and Knierim, 2012). Granule cells almost always had place fields in zero or few environments, while mossy cells generally had place fields in all or most environments

(GoodSmith et al., 2017). Using this selection criteria, 106 putative granule cells and 24 putative mossy cells were selected to be used as training data for the random forests classifier in the current study.

Although cell type labels were assigned to the training data using information about spatial firing during behavior, no spatial firing features were used for classification. The features used to classify cell types were mean firing rate, burstiness (proportion of interspike intervals that were < 6ms), and channel slope (slope of best fit line through normalized, sorted waveform peaks of the four tetrode wires) (GoodSmith et al., 2017; Neunuebel and Knierim, 2012). For classification of the cell types in mice from the current study, these features were extracted from post-behavior baseline recordings while the animal was resting or sleeping. The random forests classifier consisted of 300 individual decision trees, and two random features were used at each split. The out-of-bag error rate (an estimate of the classifier’s generalization error) was 2.3%. By applying this classifier to post-behavior sleep data recorded in the present study, we were able to identify cells as putative granule cells and mossy cells. As a measure of classification confidence, we calculated the consistency of the output of individual trees within the ensemble classifier. The random forests classifier would label a cell as a mossy cell if the majority of individual decision tree classifiers identified it as a putative mossy cell. Therefore, whether 51% or 100% of classifiers were in agreement, the overall random forest output would be the same.

For comparisons between the two cell types, we only used the most confidently classified cells (≥ 85% of individual classifier “votes”) for our primary analyses.

#### Statistics

All statistical analyses were performed in Matlab (Mathworks) and R. Data are presented as means ± SEM, and the results are considered statistically significant when p-values are less than 0.05. Wilcoxon rank-sum tests and χ^2^ tests were used to assess the difference in observed data between the two cell types. Comparisons across each session pair were made using the Kruskal-Wallis one-way ANOVA test followed by two-sided pairwise Dunn’s test with Šidák’s adjustment for post-hoc analysis. For population vector analysis, error bars represent lower and upper bounds of 95 % of the two-tailed confidence interval.

---

**Figure S1.**
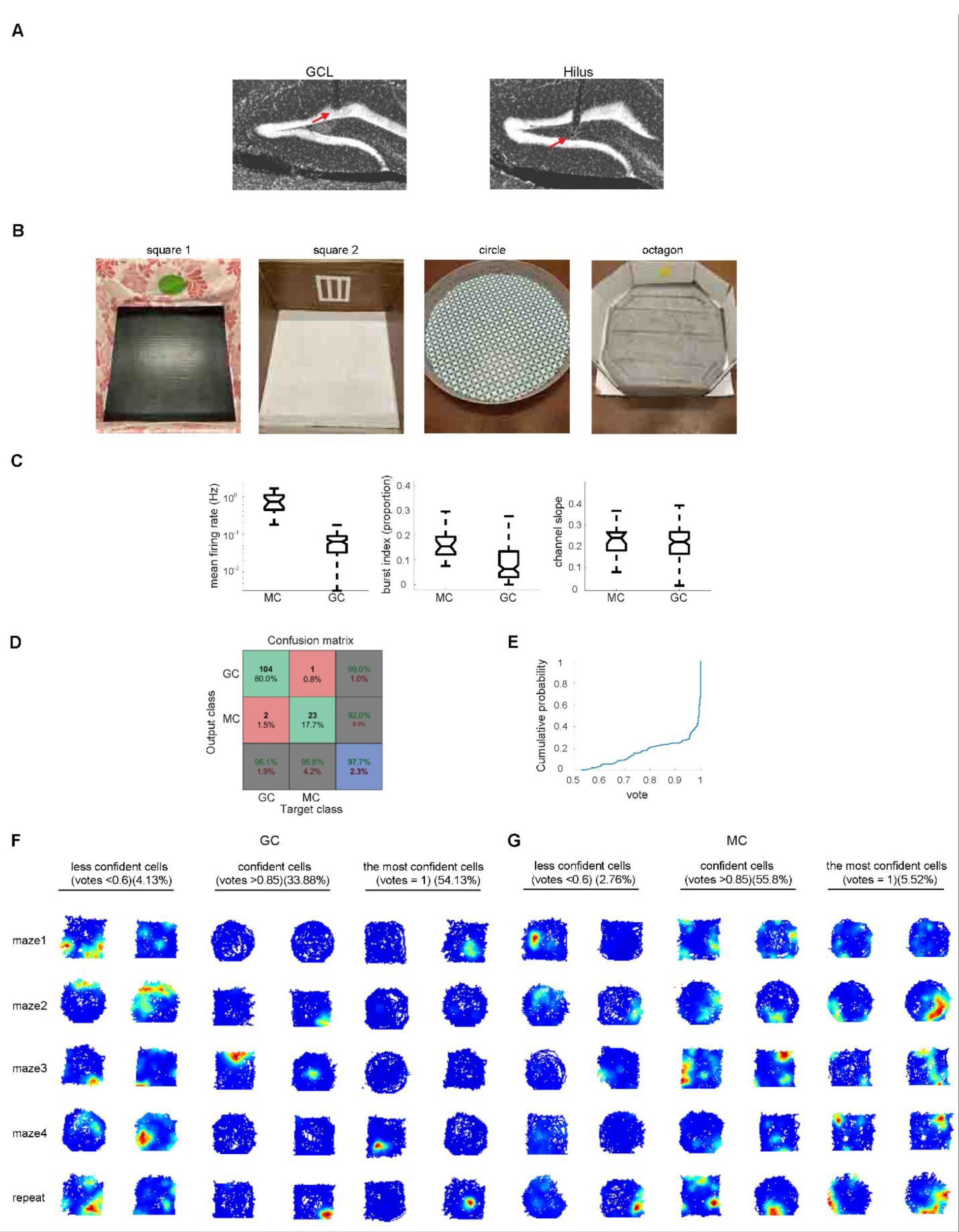
Random forests model and firing rate maps of granule cells and mossy cells based on confidence level, related to Figure 1. (A) Images of tetrode tracks terminating in the granule cell layer (GCL) and hilus. Scale bar: 200 µm. (B) Images of 4 distinct environments (two squares, circle and octagon). (C) Boxplots of features used for random forests classifier: mean firing rate (left), burst index (middle) and channel slope (right). (D) Confusion matrix for the random forests model for classification of all DG cells, comparing output class and target class from the training data. The numbers indicate number of cells from rat training data and the overall out-of-bag error rate of the classifier was 2.3%. (E) CDF of confidence level of all classified DG cells. (F) Example rate maps of granule cells (less confident cells (left), confident cells (middle) and the most confident cells (right). (G) Examples of mossy cells (less confident cells (left), confident cells (middle) and the most confident cells (right). (F-G) Each column represents recordings from a single cell over 5 sessions.

**Figure S2.**
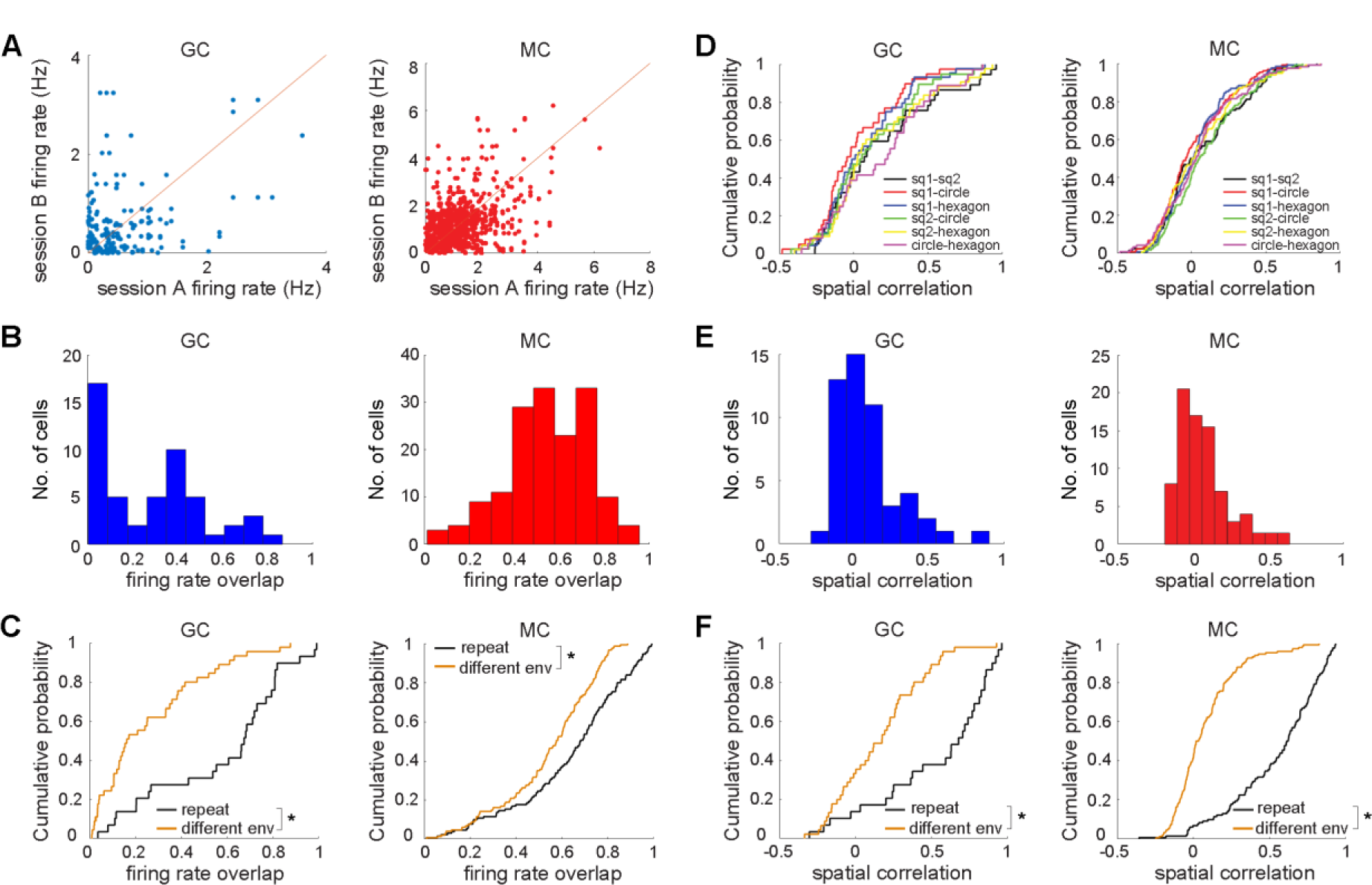
Rate overlap and spatial correlation comparison using all classified cells, without the restriction of an 85% confidence level, related to Figures 2 and 3. (A-C) Rate overlap comparison among four distinct sessions (A and B) and with a repeat session (C). Shown in (A) is a comparison of the mean firing rates from each session pair (4 environments, 3-6 pairs per cell). Shown in (B) is the histogram of firing rate overlap for granule cells (left) and mossy cells (right). Shown in (C) is the CDF of the rate overlap for granule cells (left) and mossy cells (right). CDF of repeated session (black), different environment (orange). (D-F) Spatial correlation comparison among four distinct sessions (D and E) and with a repeat session (F). Shown in (D) is the CDF of the spatial correlation for granule cells (left) and mossy cells (right) from each pair of sessions. Shown in (E) is the histogram of the spatial correlations for granule cells (left) and mossy cells (right). Shown in (F) is the CDF of the spatial correlation for granule cells (left) and mossy cells (right). CDF of repeated session (black) and different environment (orange). Statistical results in Tables S2 and S3.

**Figure S3.**
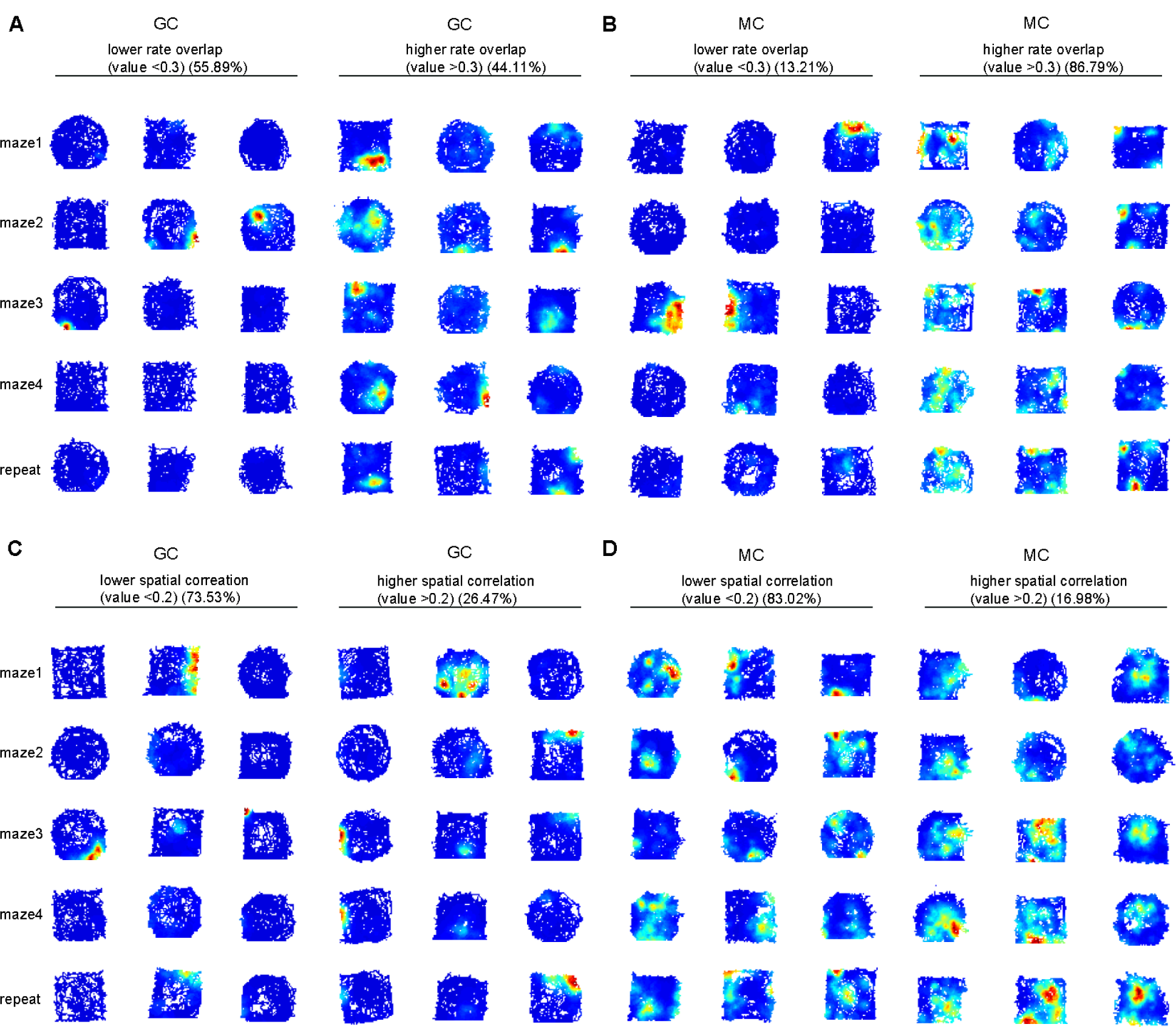
Firing rate maps of granule cells and mossy cells based on rate overlap and spatial correlation, related to Figure 2 and 3. (A) Granule cells having lower (left) and higher (right) rate overlap. (B) Mossy cells having lower (left) and higher (right) rate overlap. (C) Granule cells having lower (left) and higher (right) spatial correlation. (D) Mossy cells having lower (left) and higher (right) spatial correlation. (A-D) Each column represents recordings from a single cell over 5 sessions.

**Figure S4.**
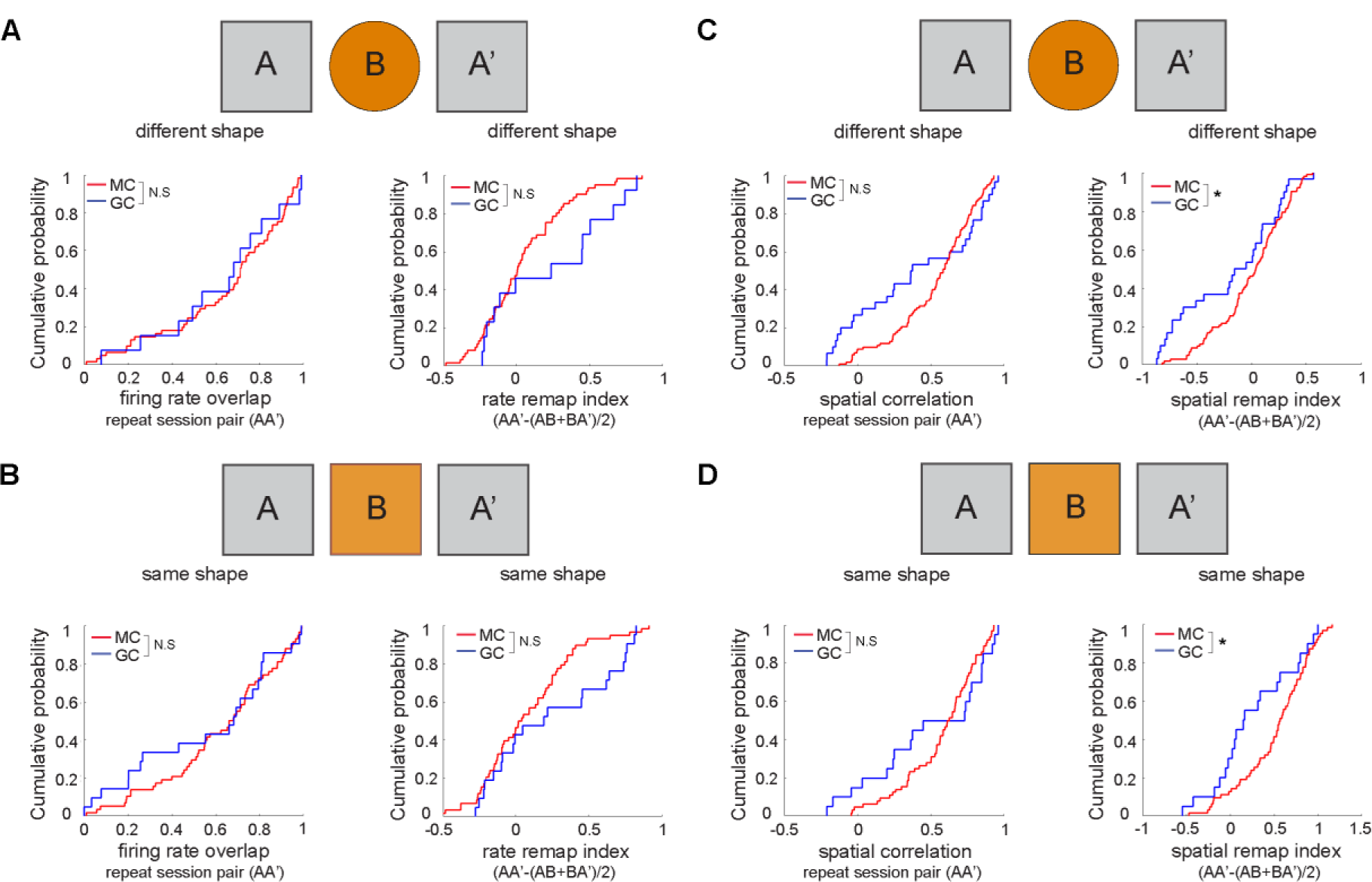
Rate and spatial remapping index using a method from a previous report (Senzai and Buzsaki 2017), related to Figure 3. Senzai and Buzsaki (2017) calculated a remapping index based on all cells that had a place field in any one of the 3 recording sessions (A, B, or A’, a repeat of session A). For reasons stated in the main text, we included only cells that had a place field in either A or A’, so as to avoid including in the analysis cells that were mostly silent or had poor place fields in both sessions, resulting in artificially low correlation values between A and A’. The comparison between A and A’ is meant to provide a baseline of firing stability between repeated exposures to the same environment, against which to compare changes in firing between different contexts. However, a cell that has poor spatial firing, or is mostly silent, in both A and A’ can be considered as firing consistently in both sessions, but it is likely to have a low correlation value nonetheless. To examine whether the difference between our results and those of Senzai and Buzsaki were due to the different inclusion criteria, we repeated our analyses using the inclusion criteria from Senzai and Buzsaki (2017). The details for statistical results are shown in Table S4. (A) CDF of rate overlap for different shapes. Left: repeat session (AA). Right: rate remapping index. (AA’-(AB+BA’)/2). Granule cells and mossy cells were not different from each other. (B) CDF of rate difference for same shape. Left: repeat session (AA). Right: rate remapping index. (AA’-(AB+BA’)/2). Granule cells and mossy cells were not different from each other. (C) CDF of spatial correlation for different shapes. Left: repeat session (AA). Right: spatial remapping index. (AA’-(AB+BA’)/2). The spatial remap index for granule cells was lower than for mossy cells (p = 0.04). (D) CDF of spatial correlation for same shape. Left: repeat session (AA). Right: spatial remapping index. (AA’-(AB+BA’)/2) A’ indicates the repeat session. Any cells with place fields in either A, B, or A’ were included for analysis. The spatial remap index for granule cells was lower than for mossy cells (p = 0.03). Thus, although the results using the two inclusion criteria are qualitatively similar (compare Figure 3 with Figure S4), the differences between granule cells and mossy cells are not significant using our criteria (Fig. 3) but they replicate the significant differences reported by Senzai and Buzsaki (2017) using their criteria. This difference may be explained in the current data by comparing the index values for the cells that were included in our analyses (Fig. 3) and the minority of cells (6/63 total mossy cells and 4/20 total granule cells in the same shape analyses; 16/102 mossy cells and 11/30 granule cells in the different shape analyses) that were included in Figure S4 but excluded from Figure 3 because they had little firing and/or poor spatial tuning in both A and A’. For the different shape analyses, the mean A-A’ correlation for granule cells meeting our inclusion criteria was 0.56, whereas the mean correlation for the 11 cells that were included using the Buzsaki criteria but excluded using our criteria was 0.12. For the same shape analyses, the mean A-A’ correlation for granule cells meeting our inclusion criteria was 0.53, whereas the mean correlation for the 4 cells that were included using the Buzsaki criteria but excluded using our criteria was 0.35. Thus, including the latter cells brought down the average of the granule cell remap index enough to make the difference statistically (and in our view artificially) significant. Although it is not clear whether using our inclusion criteria would result in the Senzai and Buzsaki data set matching our main results (Senzai and Buzsaki, personal communication), it is important to note that the differences between the studies are quantitative, not qualitative, and we believe that the inclusion criteria we used are appropriate to answer the questions at hand.

**Table S1.** Statistics for activity and remap comparisons between male and female mice, related to Figure 1 and Figure 2.

| Analysis |  | Male | Female | Test | p |
| --- | --- | --- | --- | --- | --- |
| Mean rate | GC | $1.00 \pm 0.23$ | $0.43 \pm 0.10$ | rank-sum | $z = 1.55, p = 0.12$ |
| | MC | $2.23 \pm 0.15$ | $2.05 \pm 0.26$ | rank-sum | $z = 0.89, p = 0.37$ |
| Peak rate | GC | $11.52 \pm 2.38$ | $8.64 \pm 2.07$ | rank-sum | $z = 0.52, p = 0.61$ |
| | MC | $17.84 \pm 0.89$ | $21.18 \pm 2.36$ | rank-sum | $z = -0.93, p = 0.35$ |
| Cells with place fields | GC | 23/134 | 11/78 | | $X_{(1)}^2 = 3.75, p = 0.05$ |
| | MC | 84/85 | 22/26 | | $X_{(1)}^2 = 9.34, p = 2.24 \times 10^{-3}$ |
| Rate overlap value | GC | $0.29 \pm 0.04$ | $0.20 \pm 0.07$ | rank-sum | $z = 1.25, p = 0.21$ |
| | MC | $0.56 \pm 0.02$ | $0.40 \pm 0.04$ | rank-sum | $z = -3.70, p = 2.12 \times 10^{-4}$ |
| Spatial correlation value | GC | $0.13 \pm 0.06$ | $0.06 \pm 0.05$ | rank-sum | $z = 0.37, p = 0.71$ |
| | MC | $0.05 \pm 0.02$ | $0.07 \pm 0.03$ | rank-sum | $z = -0.49, p = 0.62$ |

**Table S2.**
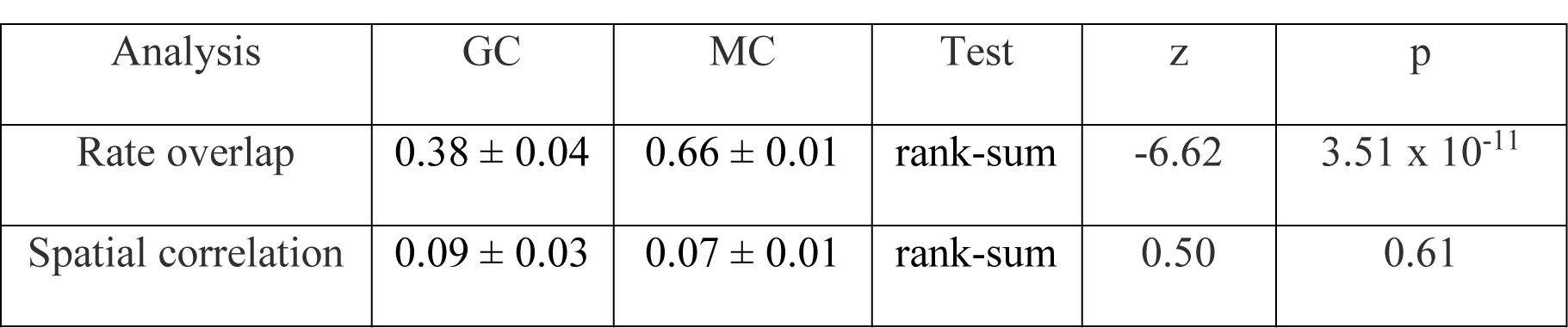
Statistics for rate overlap and spatial correlation comparison among four distinct sessions using all classified cells, related to Figure S2B and S2E.

**Table S3.** Statistics for rate overlap and spatial correlation comparison with a repeat session using all classified cells, related to Figure S2C and S2F.

| Analysis | Cell type | Repeat | Different environment | Test | z | p |
| --- | --- | --- | --- | --- | --- | --- |
| Rate overlap | GC | $0.57 \pm 0.05$ | $0.26 \pm 0.03$ | rank-sum | 4.26 | $2.02 \times 10^{-5}$ |
| | MC | $0.64 \pm 0.02$ | $0.54 \pm 0.02$ | rank-sum | 3.98 | $6.91 \times 10^{-5}$ |
| Spatial correlation | GC | $0.53 \pm 0.07$ | $0.16 \pm 0.04$ | rank-sum | 4.02 | $5.83 \times 10^{-5}$ |
| | MC | $0.55 \pm 0.03$ | $0.08 \pm 0.02$ | rank-sum | 11.25 | $2.23 \times 10^{-29}$ |

**Table S4.** Statistics for rate and spatial remapping indices using a method from previous report (Senzai and Buszaki 2017), related to **Figure S4**.

|  |  | Test | Rate remap index |  |  |  | Spatial remap index |  |  |  |
| --- | --- | --- | --- | --- | --- | --- | --- | --- | --- | --- |
|  |  |  | GC | MC | z | p | GC | MC | z | p |
| **Different shape** | AA' | rank-sum | $0.64 \pm 0.08$ | $0.66 \pm 0.03$ | 0.28 | 0.78 | $0.40 \pm 0.08$ | $0.53 \pm 0.03$ | -1.11 | 0.27 |
| | Remap index | rank-sum | $0.23 \pm 0.11$ | $0.04 \pm 0.04$ | -1.29 | 0.20 | $0.31 \pm 0.08$ | $0.49 \pm 0.03$ | -2.04 | 0.04 |
| **Same shape** | AA' | rank-sum | $0.56 \pm 0.07$ | $0.62 \pm 0.03$ | 0.53 | 0.60 | $0.49 \pm 0.09$ | $0.58 \pm 0.03$ | -0.28 | 0.78 |
| | Remap index | rank-sum | $0.25 \pm 0.09$ | $0.08 \pm 0.04$ | -1.33 | 0.18 | $0.26 \pm 0.10$ | $0.52 \pm 0.05$ | -2.11 | 0.03 |

